# Tools, strains, and strategies to effectively conduct anaerobic and aerobic transcriptional reporter screens and assays in *Staphylococcus aureus*

**DOI:** 10.1101/2021.06.11.448164

**Authors:** Erin E. Price, Paulami Rudra, Javiera Norambuena, Franklin Román-Rodríguez, Jeffrey M. Boyd

## Abstract

Transcriptional reporters are reliable and time-tested tools to study gene regulation. In *Staphylococcus aureus*, β-galactosidase (*lacZ*)-based genetic screens are not widely used because of the necessity of selectable markers for strain construction and the production of staphyloxanthin pigment which obfuscates results. We describe a series of vectors that allow for markerless insertion of codon-optimized *lacZ*-based transcriptional reporters. The vectors encode for different ribosomal binding sites allowing for tailored *lacZ* expression. A Δ*crtM::kanR* deletion insertion mutant was constructed that prevents the synthesis of staphyloxanthin, thereby permitting blue-white screening without the interference of carotenoid production. We demonstrate the utility of these vectors to monitor aerobic and anaerobic transcriptional activity. For the latter, we describe the use of a ferrocyanide-ferricyanide redox system (Fe(CN)_6_^3–/4–^) permitting blue-white screening in the absence of oxygen. We also describe additional reporter systems and methods for monitoring transcriptional activity during anaerobic culture including a FAD-binding fluorescent protein (*EcFbFP*), alpha-hemolysin (*hla*), or lipase (*geh*). The systems and methods described are compatible with vectors utilized to create and screen high-density transposon mutant libraries.

**Importance:** *Staphylococcus aureus* is a human pathogen and a leading cause of infectious disease-related illness and death worldwide. For *S. aureus* to successfully colonize and invade host tissues, it must tightly control the expression of genes encoding for virulence factors. Oxygen tension varies greatly at infection sites and many abscesses are devoid of oxygen. In this study, we have developed novel tools and methods to study how and when *S. aureus* alters transcription of genes. A key advantage to these methods and tools is that they can be utilized in the presence and absence of oxygen. A better understanding of anaerobic gene expression in *S. aureus* will provide important insight into the regulation of genes in low oxygen environments.

## Introduction

*Staphylococcus aureus* is a Gram-positive pathogen causing morbidity and mortality worldwide. *S. aureus* produces numerous virulence factors which contribute to bacterial pathogenesis. Understanding how and when *S. aureus* alters transcription of genes encoding virulence factors is key to understanding pathogenesis.

β-galactosidase (*lacZ*) assays have been widely used to study the function of bacterial gene regulatory elements by allowing for quantification of promoter activity on gene expression. The *Escherichia coli* LacZ (120 kDa, 1024 amino acids) has β-galactosidase activity, which catalyzes the hydrolysis of β-galactosides into monosaccharides. β-galactosidase has high stability, is resistant to proteolytic degradation, and does not significantly degrade or bleach (as occurs with fluorescent reporters), which promotes successful usage of β-galactosidase as a transcriptional reporter (1, 2). Miller described a standardized protocol for measuring β-galactosidase activity using the synthetic substrate *o*-nitrophenyl-β-d-galactoside (ONPG) (3). The hydrolysis of ONPG leads to production of the colored compound *o*-nitrophenol (ONP), which can be measured spectrophotometrically; moreover, monitoring the hydrolysis of ONPG is fast, inexpensive, consistent, and sensitive. The compound X-Gal (5-Bromo-4-chloro-3-indolyl β-D-galactopyranoside) is hydrolyzed by LacZ to produce 5-bromo-4-chloro-3-hydroxyindole, which can be oxidized to form 5,5-dibromo-4,4’dichloro-indigo, an insoluble indigo precipitate (4). Dimerization of 5-bromo-4-chloro-3-hydroxyindole is widely used to detect β-galactosidase activity on solid media during aerobic culture where dioxygen serves as the oxidant.

*lacZ*-based technologies have been used in *S. aureus* previously. O’Neill *et al*. developed a β-galactosidase leakage assay to assess the ability of molecules to cause membrane damage (5). To this end, they created a *S. aureus* strain carrying the *E. coli lacZ* gene under the transcriptional control of a strong staphylococcal promoter (*cap1A*). The strains were exposed to various membrane-damaging substances and leakage was detected by monitoring the activity of β-galactosidase in the cell-free supernatant using a fluorescence assay with 4-methylumbelliferyl-β-d-galactoside as a substrate. Similarly, Ranjit *et al*. used β-galactosidase leakage assays to examine the role of disulfide bond formation in cell lysis and oligomerization of a membrane-associated holin protein CidA (6). Baum *et al*. constructed *S. aureus* strains in which chromosomal insertions contained *lacZ* under the transcriptional control of the *msrA1* or *msrB* promoters (7). These strains were used to monitor *lacZ* expression after generating additional chromosomal mutations and after the addition of the cell wall-active antibiotic oxacillin.

In addition to quantitative β-galactosidase activity assays, *lacZ-*based colorimetric screening experiments have also been previously employed in *S. aureus*. Nielsen *et al*. developed an assay to screen compounds that influence virulence factor production in *S. aureus* using transcriptional *lacZ* reporters fused to the promoter sequences of virulence factor genes *hla, rnaIII*, and *spa* (8). These strains were placed as top agar overlays in medium containing X-gal substrate, and cell-free fungal lysates containing potential compound(s) of interest were spotted upon the overlay. Blue-white color development in the agar overlay was monitored to indicate changes in promoter activity. More recently, Bojer *et al*. used a similar approach to investigate the effects of antimicrobial peptides on virulence gene expression (9). Another work by Ding *et al*. demonstrated the influence of a citrate responsive catabolite control protein E (CcpE) on the promoter activity of aconitase gene (the second enzyme of the tricarboxylic acid cycle) *citB* (10). They created transcriptionally fused *citB-lacZ* and found that not only was the promoter activity of *citB* considerably reduced in the Δ*ccpE* mutant than in the wild-type strain, but also the mutation of box-I sequence in *citB* promoter completely abolished the promoter activity. LacZ-based technologies have not been widely applied to non-biased genetic screens in *S. aureus*.

To our knowledge, anaerobic monitoring of *lacZ* expression using X-Gal has not been utilized because the dimerization of the reaction intermediate monomers (5-bromo-4-chloro-3-hydroxyindol) requires oxidation to form blue precipitate (5,5’-dibromo-4,4’-dichloro-indigo). Other reporter systems, including luciferase- and GFP-based fluorescent proteins, also require oxygen for signal output. Recently, a new class of oxygen-independent flavin mononucleotide-based fluorescent proteins (FbFPs) have been characterized (11, 12). Drepper *et al*. engineered a set of FbFPs that are derivatives of bacterial blue-light receptors from *Bacillus subtilis* and *Pseudomonas putida* (12). These proteins were used to generate fluorescent reporter systems that are functional under both aerobic and anaerobic conditions in *E. coli* (EcFbFP).

Alpha-hemolysin (alpha-toxin), encoded by *hla*, is a prototypic β-barrel toxin and one of the key virulence factors of *S. aureus*. Upon secretion, it forms a pore in the membranes of target host red blood cells, resulting in cell lysis (13). The *S. aureus* genome encodes for several secreted lipase enzymes, which serve to break down host-derived lipids into free fatty acids for nutrient acquisition (14). Of these, the glycerol ester hydrolase lipase is encoded by the *geh* locus and is specific for long-chain fatty acids (15, 16).

Here, we describe vectors that allow for markerless transcriptional reporters utilizing *lacZ, EcFbFP, hla*, or *geh* expression to monitor promoter activity. The vectors allow for expression to be driven by different ribosome binding sequences (RBS) of varied strength. Importantly, when the vectors are resolved after making mutants, they do not leave behind genetic determinants that provide antibiotic resistance, and therefore can be used for additional genetic manipulations including the generation of transposon mutant libraries. We also describe a *crtM::kan* deletion insertion mutation that prevents staphyloxanthin production and aids in mutant identification during blue-white screening. We outline methods to use the vectors for both aerobic and anaerobic screening.

## Materials and Methods

### Bacterial strains and culture conditions

Tryptic soy broth (TSB) was purchased from VWR. X-gal was purchased from VWR and 3,4-Cyclohexeneoesculetin-B-D-galactopyranoside sodium salt (S-gal) was purchased from Sigma-Aldrich. For solid medium (TSA), TSB was supplemented with 1.5% agar. For aerobic spotting assays, individual strains were grown in 5 ml of TSB in 30-ml culture tubes and shaken at 220 rpm at 37°C to an optical density at 600 nm (OD_600_) of 1. Strains were serially diluted and 5 µl were spotted as 10-fold dilutions on TSA plates containing varying concentrations of X-gal or S-gal. For plates containing S-gal, the agar was also supplemented with ferric ammonium citrate (Sigma-Aldrich) at a concentration of 62.5 µg ml^-1^. For anaerobic spotting experiments, plates were incubated at 37°C within a COY anaerobic chamber for 36 hours. Overlay plates were sprayed carefully in the chamber with X-gal (25 mg ml^-1^ prepared in DMSO) supplemented with tetracycline (3.3 mg ml^-1^) until the agar surface was completely covered. Sprayed plates were removed from the chamber and exposed to oxygen. Plates were dried in a fume hood and were developed for an hour.

When selecting for plasmids, episomes, or chromosomal insertions, antibiotics were added at the following final concentrations: 150 μg ml^-1^ ampicillin, 30 μg ml^-1^ chloramphenicol (Cm), 10 μg ml^-1^ erythromycin (Erm), 50 μg ml^-1^ kanamycin (Kan) or 3 μg ml^-1^ tetracycline (Tet).

### Plasmid and strain construction

All transductions were conducted using bacteriophage 80α (17). All bacterial strains were PCR verified before use. Plasmids were sequenced at Genewiz (South Plainfield, NJ). Synthetic DNA was synthesized by Twist Biosciences (San Francisco, CA) or Integrated DNA Technologies (Coralville, IA). DNA primers were purchased from Integrated DNA Technologies (Coralville, IA). Phusion DNA polymerase was purchased from New England Biolabs. The *lacZ* and *EcFbFP* were codon-optimized for expression in *S. aureus* using the online Integrated DNA Technologies codon optimization tool.

Yeast homologous recombination was used to construct plasmids as previously described (18, 19). To begin, portions of DNA were synthesized containing: (i) a 3’ portion homologous to an upstream portion of *geh*, (ii) a polylinker, (iii) the *saeP1* or *suf* promoter, (iv) KpnI, NheI, MluI and SalI restriction sites, (v) a *sodM, sarA, hld*, or *TIR* ribosomal binding site, and (vi) the 5’ portion of the codon-optimized *lacZ*. The sequences of the synthesized DNA constructs used for construction of plasmids are listed in **Table S1**. The sequences of the DNA primers utilized to generate PCR amplicons are listed in **Table S2**. The yeast cloning cassette and *lacZ* sequences were amplified using pJB38_Δ*copBL* and the pJB185 as templates, respectively (20). The plasmids that were created using yeast recombinational cloning along with the DNA primers and DNA templates used to generate the amplicons are listed in **Table S3**. The amplicons were combined with EcoRI-digested pJB38 and transformed into *Saccharomyces cerevisiae* strain FY2. *S. cerevisiae* colonies containing the plasmid of interest were identified by colony PCR and further propagated. Plasmids were recovered from yeast and electroporated into *E. coli* PX5 (Protein Express) cells selecting for Amp resistance. The plasmids were then transformed into *S. aureus* RN4220 and selected for Cm resistance. The vectors were transduced into JMB1100 and the integrates were constructed as previously described (21).

In order to create Δ*crtM::kanR* deletion strain, approximately 500 base pairs upstream and downstream of the *crtM* gene (SAUSA300_2499) were PCR amplified using JMB1100 or JMB2122 (*kanR*) chromosomal DNA as a template, and the following primer pairs, respectively: YCC_crtM_for and kanR_up_crtM_rev; up kanR_crtM_for and down_kanR_crtM_rev; kanR_down_crtM_for and pJB38_crtM_rev. pJB38_*rseE::tet* was digested with MluI and NheI and gel purified. The vector and amplicons were combined and transduced into *S. cerevisiae* strain FY2 resulting in pJB38_Δ*crtM::kanR*. The plasmid was recovered using *E. coli* PX5 and transformed into *S. aureus* strain RN4220. The mutant was created in JMB 1100 resulting in strain JMB9964 and then the *crtM::kanR* allele was transduced into strains of interest.

To generate the pJB38_*saeP1*_*sarA* RBS_*EcFbFP* vector, the pJB38_*saeP1*_*TIR* RBS_*lacZ* vector was digested with MluI and PstI. MluI cuts upstream of the *lacZ*. PstI cuts downstream of the *geh* downstream fragment. The vector backbone was gel purified and combined with the amplicons generated using the following primer pairs: gehpJB38 pstI and EcFbFP*geh*for; P1TIR EcFbFP for and EcFbFP*geh*rev. Chromosomal DNA and the synthesized EcFbFP fragment were used as templates for PCR, respectively. To generate the pCM28_*saeP1*_*sarA* RBS_*EcFbFP* vector, pCM28 was digested with BamHI and SalI. The gel purified vector backbone was combined with amplicons generated using pJB38_*saeP1*_*sarA* RBS_*EcFbFP* as a PCR template and the following primer pairs: pCM28_YCC_for and YCC_P1_rev; YCC_P1_for and EcFbFP_pCM28_3. Note that the upstream and downstream primers did not contain the BamHI and SalI restriction sites found in pCM28, yielding a plasmid lacking these sites.

To create the pJB38_*saeP1*_*TIR* RBS_*hla* pJB38_*saeP1*_*TIR* RBS_*geh* vectors, pJB38 was linearized with EcoRI. The native *hla* or *geh* promoters and RBS were replaced with the *saeP1* promoter and either the *sodM* or *TIR* RBS. This was flanked by approximately 500 base pairs of chromosomal DNA from upstream of the promoter and 500 base pairs of downstream chromosomal DNA that initiates that the translational start site. The vectors integrate at the native *hla* or *geh* loci and replace the native promoter with the *saeP1* promoter and the selected RBS.

Several additional plasmids were created using restriction enzyme-based cloning. These plasmids, as well as the primers, vector backbones, and restriction enzymes that were utilized to create them, are listed in **Table S4**. Representative vector sequences and maps are shown in **Table S5**.

### Quantitative β-galactosidase assay

The bacterial strains were grown to an OD_600_ of approximately 1. Cell culture (1 ml) was pelleted by centrifugation. Cell pellets were resuspended in 1.2 ml of Z-buffer (60 mM Na_2_HPO_4_, 40 mM NaH_2_PO_4_, 10 mM KCl, 1 mM MgSO_4_, 50 mM β-mercaptoethanol, pH 7.0), in 2-ml screw cap tubes containing 0.1 mm silica glass beads (MP Biomedicals). Cells were lysed by bead beating (3 cycles, 40s each, 6.0 m/s) using a FastPrep homogenizer (MP Biomedicals). Material was centrifuged at 13,000×g for 2 min to remove unlysed cells and insoluble debris. Supernatant lysate (20 µl) was added to 680 µl of Z-buffer. 140 µl of ONPG (4 mg ml^-1^ (w/v)) was added to the samples. The reactions were quenched by adding 200 µl stop solution (1 M Na_2_CO_3_) as soon as the samples turned light yellow and the reaction time was recorded. The A_420_ of the samples were measured using a UV spectrophotometer (Beckman Coulter DU^®^ 530 Life Science UV/Vis Spectrophotometer). The corresponding protein concentrations of the samples were measured using a Bradford assay as previously described [15]. The Modified Miller Units (specific activity) were calculated as following:

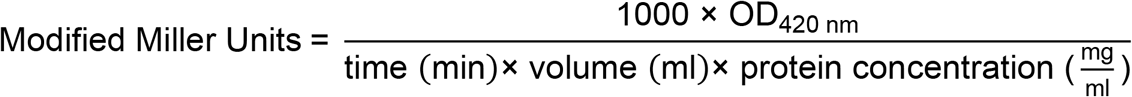

For anaerobic β-galactosidase assays, the reporter strains were grown aerobically to an OD_600_ of ∼1, then transferred to a 37°C incubator in an anaerobic chamber and grown statically overnight. The cells were pelleted inside the anaerobic chamber by centrifugation, resuspended in 1.2 ml of Z-buffer, and transferred to glass-bead-containing screw cap tubes prior to removal from the anaerobic chamber. The remaining procedure after cell lysis was carried out as described above.

### Transposon library construction

Transposon mutant libraries were constructed in the *saeP1*_*sarA* RBS_*lacZ* (JMB 9709) reporter strain as previously described by Grosser *et al*. (22, 23). Briefly, the plasmid pMG020 (encoding transposase) was freshly transformed into RN4220 and incubated on TSA Tet (10 µg ml^-1^) at 30°C. Single colonies were selected and grown in TSB Tet (10 µg ml^-1^) at 30 °C and lysates were generated. Reporter strains carrying pBursa were transduced with pMG020 and selected on TSA Cm-Tet plates at 30 °C. Cells grown from individual colonies struck on TSA Cm-Tet plates were diluted and suspended in 200 µl PBS buffer, and 15 µl aliquots were spread onto TSA plates containing 10 µg ml^-1^ Erm, then incubated at 42°C for 24 hours to allow for transposition. In total, colonies from 22 large petri plates (containing approximately 3,000 colonies each) were pooled using TSB 10 µg ml^-1^ Erm supplemented with 25% glycerol. Aliquots were thoroughly mixed by vortexing and combined into a single pool of transposon mutants. Aliquots (1 ml each) were then frozen and stored at -80°C. Aliquots of the transposon libraries were plated on TSA Erm plates containing 50 µg ml^-1^ X-gal to select single colony mutants with altered *lacZ* expression. Mutants with altered *lacZ* expression were reconstructed by transforming the Tn lesion back into the parent followed by qualitative and quantitative β-galactosidase assays.

### Mapping the locations of chromosomal insertions

The genomic locations of transposon insertions were mapped as previously described, with slight modifications (22, 23). Briefly, genomic DNA was isolated from the colonies which displayed a variation in X-gal hydrolysis using a Lucigen Gram-positive DNA purification kit. Chromosomal DNA (1 µg) was digested with 10 units (1 µl) of restriction enzyme AciI (New England Biolabs (NEB)) for one hour at 37°C, then heat inactivated at 65°C for 30 minutes and ligated using 1 µl Quick Ligase (NEB) at room temperature for 15 minutes. PCR reactions were then performed in a final volume of 50 µl containing the ligated DNA, Phusion polymerase and the Tn-Buster and Martn-ermR DNA primers. DNA was amplified using a three-step PCR cycle: denaturation (98°C for 30s), annealing (50°C for 30s), and elongation (72°C for 2 min), repeated 25 times. The PCR products were separated on a 1% agarose gel, DNA bands were gel extracted (Qiagen) and submitted for Sanger sequencing using either the Tn-Buster or Martn-ermR primers.

### Examining expression of EcFbFP reporter constructs

Fluorescence measurements of whole-cell liquid cultures were carried out photometrically on a Variskan Lux plate reader (Thermo Scientific). Aliquots of cell cultures (1 ml) grown under aerobic or anaerobic conditions were pelleted by centrifugation and resuspended in 1 ml of PBS. Samples of 200 µl were used for quantification of the fluorescence intensity in a black 96-well plate at room temperature. Measurements were taken with an excitation wavelength of 450 nm, an emission wavelength of 495 nm, and a 12 nm path length. Triplicate samples from each strain were averaged and normalized against cell density.

### Hemolysin and lipase activity assays

For plate-based hemolysis and lipase assays, overnight cultures grown in TSB were diluted and grown to an optical density of 1 (A_600_). For hemolysis assays, two microliters of the cell suspension were spotted onto TSA plates containing 5% defibrinated rabbit blood (HemoStat Laboratories). For lipase assays, two microliters of cell suspension were spotted onto lipase activity plates (1% peptone, 85 mM NaCl, 8.8 mM CaCl_2_, 1.5% agar) containing 1% Tween-80 (VWR). Plates were incubated at 37°C aerobically or in a COY anaerobic chamber until halos surrounding the spotted cells appeared.

Quantitative hemolysis assays were performed as previously described with slight modifications (24). Briefly, cultures were incubated at 37°C with shaking at 220 rpm for 18 h. For anaerobic samples, strains were grown aerobically to an OD_600_ of ∼1 and then transferred to a 37°C incubator in an anaerobic chamber and grown statically overnight. The cultures were diluted with TSB to equalize the OD_600_ to ∼0.05, pelleted by centrifugation, and sterilized through a 0.2-µm filter. Samples (100 µl) were incubated at 37°C with a 3% solution of PBS-washed rabbit blood cells in a BioTek Epoch 2 microplate reader, and heme content was assessed with absorbance measurements at 450 nm taken every 4 min for 2 hours. Biological triplicates were assayed in duplicate and averaged.

## Results

### Creation of *S. aureus lacZ* transcriptional reporters

We envisioned a series of plasmids containing transcriptional reporters that could be used to generate markerless *S. aureus* chromosomal insertions. We chose to use the pJB38 vector as a backbone, which encodes for ampicillin and chloramphenicol resistance in *E. coli* and *S. aureus*, respectively [18]. It also encodes for a temperature sensitive origin of replication in *S. aureus* and for an anhydrotetracycline inducible antisense RNA to the *S. aureus secY* allowing for inducible counter selection.

We created a series of four vectors (**Figure S1**) that integrate into the *geh* locus (SAUSA300_0320), which is a commonly used as an episome integration site (25). The vectors have one of four ribosomal binding sites (RBS) that drive expression of *Escherichia coli lacZ* that was codon-optimized for *S. aureus* expression: *hld, sar, TIR*, and *sodM* (**Figure 1A)** (20). Altering the ribosomal binding site can alter gene expression in *S. aureus* (26). The *TIR* RBS was recently described to drive a constitutively high level of expression (27, 28). The vectors contain the yeast two-micron origin of replication and *URE3* allowing for the selection for uracil prototrophs using *S. cerevisiae* strain FY2, thereby permitting yeast recombinational cloning. The vectors contain a polylinker region upstream of the promoter, as well as KpnI and MluI restriction sites downstream of the promoter, yet upstream of the individual RBS (**Table S5**). These restriction sites allow for the interchanging of promoter sequences, while simultaneously preserving the location and presence of the RBS. Importantly, once the vector backbone is excised from the *S. aureus* genome, the integrated elements do not leave a gene encoding for an antibiotic resistance. This allows for strains containing the chromosomal-resolved transcriptional reporter to be used for further genetic manipulation.

**Figure 1.**
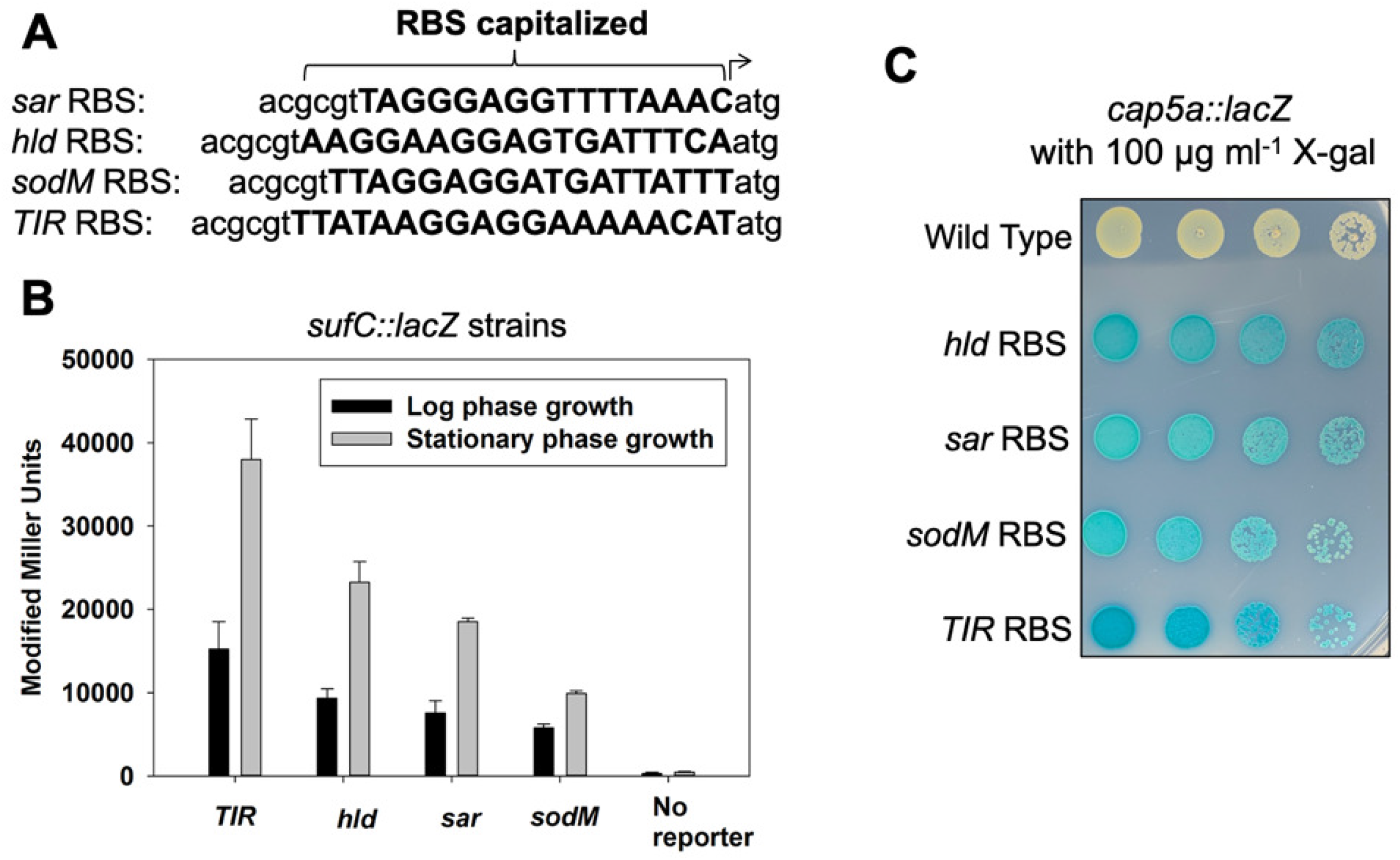
The ribosomal binding site alters *lacZ* expression. **(A)** The DNA sequences of the four ribosomal binding sites (RBS) utilized to generate the transcriptional reporters. **(B)** β-galactosidase activity in the wild type (JMB 1100: no *lacZ*), *geh::suf _TIR* RBS*_lacZ* (JMB 9741), *suf _sodM* RBS*_lacZ* (JMB 9739), *suf _sarA* RBS*_lacZ* (JMB 9740), *suf _hld* RBS*_lacZ* (JMB 9738) strains during logarithmic or stationary growth phases. **(C)** Overnight cultures of the wild type (JMB 1100), *cap5a_TIR* RBS*_lacZ* (JMB 9765), *cap5a_sodM* RBS*_lacZ* (JMB 9764), *cap5a _sarA* RBS*_lacZ* (JMB 9754), and *cap5a _hld* RBS*_lacZ* (JMB 9777) strains were serial diluted by 10-fold dilutions and 5 μl of each strain was spotted on TSA containing 100 mg ml^-1^ X-gal. The data displayed in Panel B are the average of biological triplicates with the standard deviations shown. A representative image is shown in panel C.

We first constructed a series of strains containing the promoter for the *sufCDSUB* operon driving expression of *lacZ* (29). Each of the four strains contained a different RBS sequence driving *lacZ* expression. β-galactosidase activity was quantified after liquid growth, as well as visualized on solid media. The different *suf::lacZ* strains had varied β-galactosidase activity after liquid growth resulting in the following expression pattern: *TIR*>*hld*>*sar*>*sodM* (**Figure 1B**). The alterations in RBS-driven *lacZ* expression were independent of growth phase. As predicted, the wild-type strain (JMB 1100) did not display significant β-galactosidase activity. We could not visually distinguish a difference between these reporter strains when they were spot plated on solid medium containing X-gal (data not shown).

We also examined *lacZ* expression driven by the *cap5a* promoter on solid medium containing X-gal. In this case, we could visualize variation in intensity of blue color of the *cap5a::lacZ* strains, which displayed a pattern of indole formation that was consistent with the liquid assay results: *TIR>hld>sar/sodM* (**Figure 1C**).

### Examining the effect of chromosomal manipulations on transcriptional reporter activity

SaeRS is a two-component regulatory system in *S. aureus* (30). SaeR and SaeS are the response regulator and the histidine kinase, respectively. SaeS also interacts with SaeP, which stimulates the phosphatase activity of SaeS (31). The *S. aureus sae* locus is comprised of four open reading frames (*saePQRS*), which have two promoters. The first promoter, denoted P1, lies upstream of *saeP* and is responsive to the phosphorylation status of SaeR (31).

We created a strain containing the P1 promoter driving *lacZ* expression (*geh*::*P1 sar* RBS_*lacZ*; *saeP1::lacZ*). We transduced this strain with *saeR::Tn* and *saeP::Tn* insertional inactivation mutations and examined *lacZ* expression on solid medium by spot plating the *saeP1::lacZ* strains with varying concentrations of X-gal. The strains displayed a concomitant increase in indole formation as a function of X-gal concentration (**Figure 2A**). The strain containing the *saeR::Tn* mutant displayed a visual decrease in indole formation (**Figure 2A**) when compared to the parent strain. The strain containing the *saeP::Tn* mutation behaved similarly to the parent, suggesting that SaeP was inactive under the growth conditions utilized. These data suggest that the concentration of X-gal could be varied to be effectively used to examine both activators and repressors of a locus.

**Figure 2.**
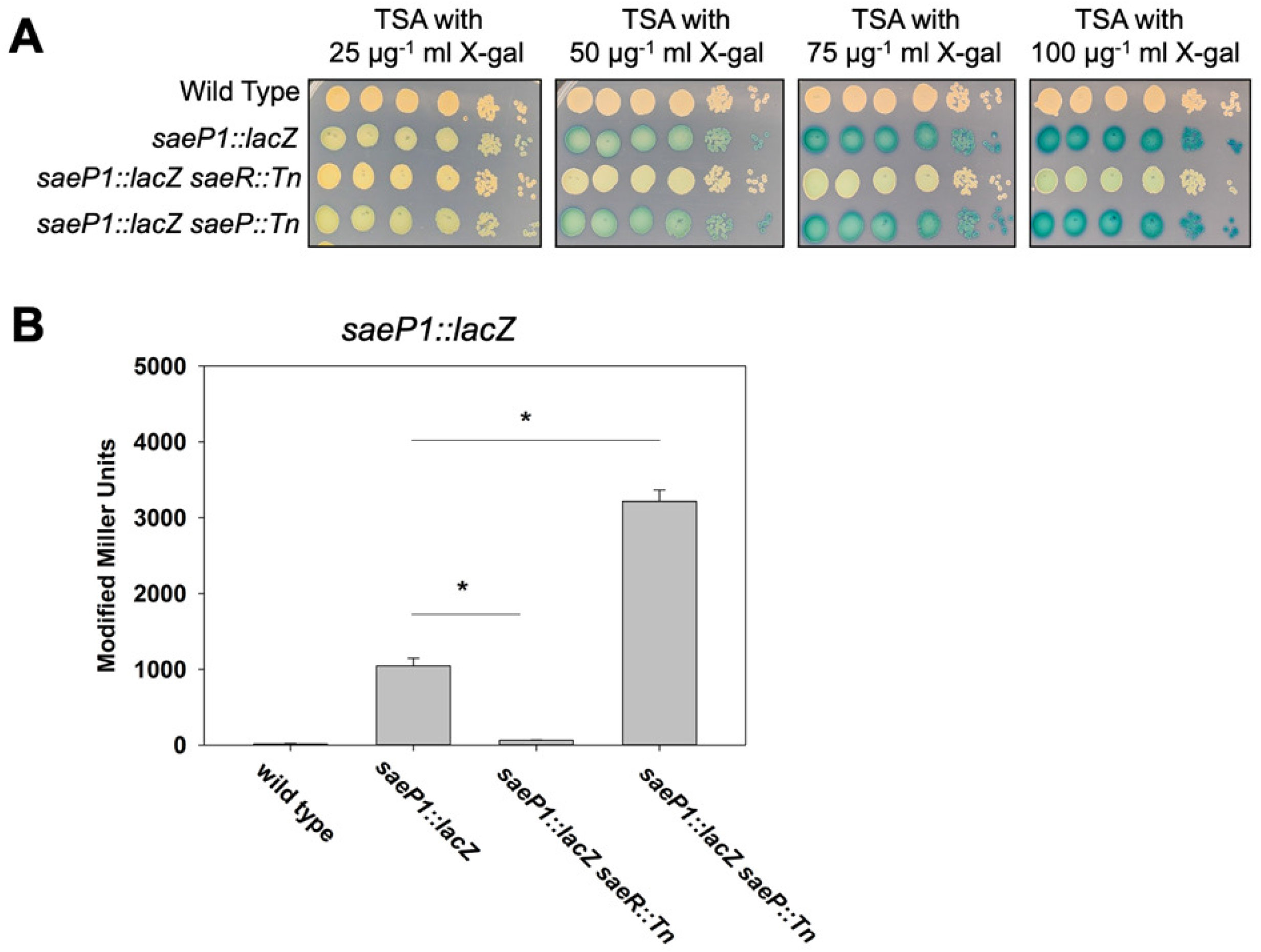
Directed chromosomal manipulations alter *saeP1::lacZ* expression. **(A)** Overnight cultures of the wild type (JMB 1100), *saeP1::lacZ* (JMB 9709), *saeP1::lacZ saeR::Tn* (JMB 9727), *saeP1::lacZ saeP::Tn* (JMB 9728) strains were serial diluted and spot plated on TSA with various concentrations of X-gal. **(B)** Quantitative β-galactosidase assays using the strains examined in Panel A. All strains utilized the *sarA* RBS to drive *lacZ* expression. Representative images are displayed in panel A. The data depicted in panel B represent the average of biological triplicates with the standard deviations shown.

We analyzed β-galactosidase expression after aerobic liquid growth. As expected, the *saeR::Tn* mutant was required to activate transcription of *saeP1::lacZ*. The *saeP::Tn* mutant had increased β-galactosidase activity when compared to the parent strain, suggesting the SaeP was stimulating phosphatase activity under this growth condition.

### Generation of Δ*crtM::kan* allele to improve the resolution of blue-white screening

We built a mariner-based transposon mutant library in a *saeP1::lacZ* (*sar_*RBS) reporter strain and screened the library for strains with altered P1 promoter activity. Several strains were isolated that visually appeared to have altered indole formation after aerobic growth. When the strains were plated on TSA medium that did not contain X-gal, they had no pigmentation (**Figure 3A**). The pigment staphyloxanthin provides most *S. aureus* strains with a characteristic golden color (32). We determined the chromosomal location of one of the insertions that generated a non-pigmented *S. aureus* strain and found that it was in the *crtP* gene locus (SAUSA300_2501) (**Figure 3A**). CrtP is one of the enzymes required for staphyloxanthin biosynthesis (33). We determined β-galactosidase activity in the parent and the *crtP* mutant after liquid growth and found that there was not a significant difference in either logarithmic or stationary phases of growth (**Figure 3B**).

**Figure 3.**
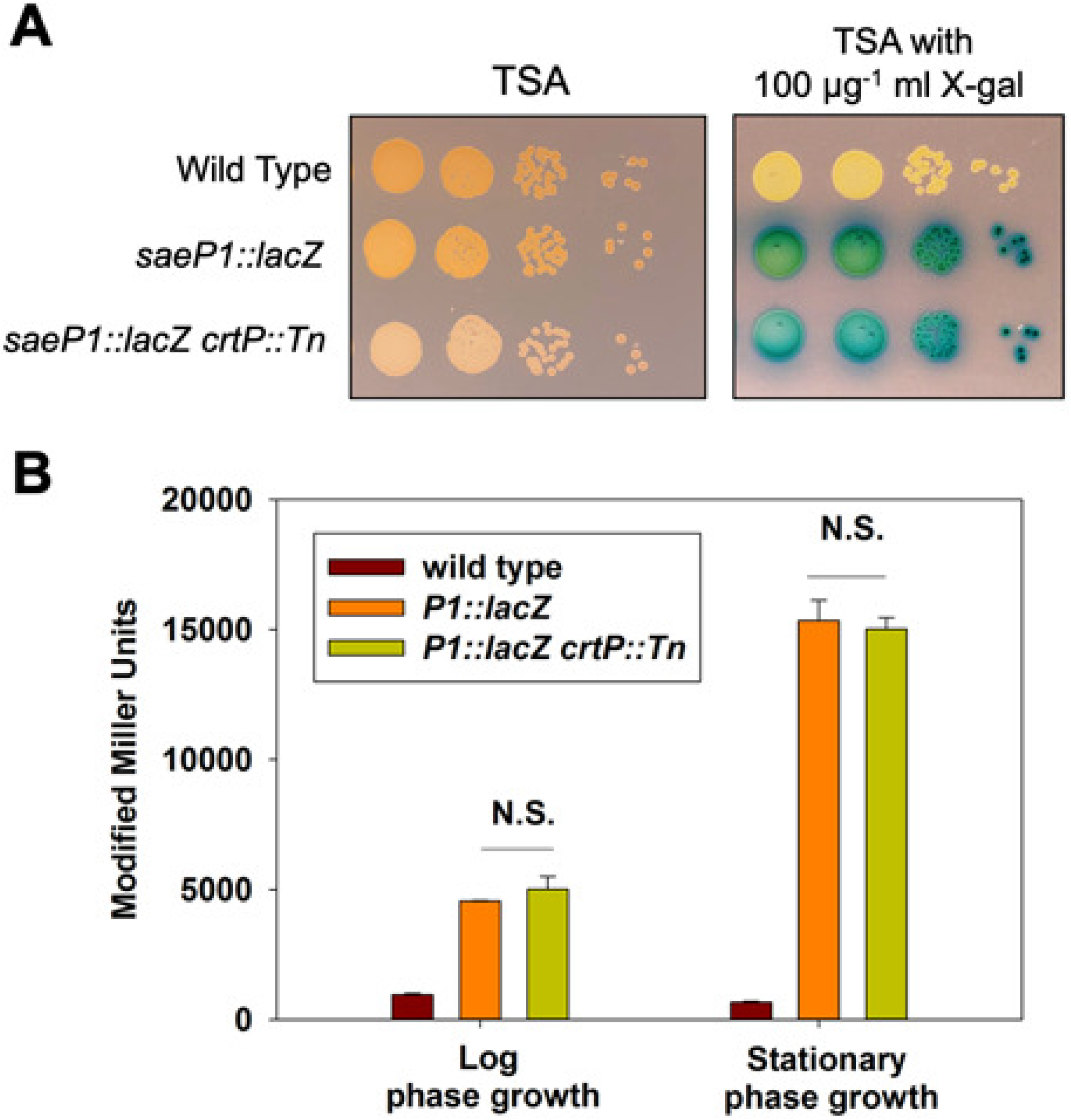
Staphyloxanthin production obfuscates visualizing indole formation. **(A)** Overnight cultures of the wild type (JMB 1100, no *lacZ*), *saeP1::lacZ* (JMB 9740), and *saeP1::lacZ crtP::Tn* (JMB 9832) strains were serial diluted and spot plated on TSA with and without 100 μg ml^-1^ X-gal. **(B)** β-galactosidase activity in the wild type (JMB 1100), *saeP1::lacZ* (JMB 9709), and *saeP1::lacZ crtP::Tn* (JMB 9832) strains during logarithmic or stationary growth phases. All strains utilized the *sarA* RBS to drive *lacZ* expression. A representative photo is displayed in panel A. The data displayed in panel B represent the average of biological triplicates with the standard deviations shown.

We hypothesized that staphyloxathin accumulation was interfering with the intensity of blue color and complicating our screening process. To facilitate better resolution for solid media *lacZ*-based blue-white screening with *S. aureus*, we constructed a strain that lacked the ability to produce staphyloxanthin. CrtM catalyzes the first committed step in staphyloxanthin production (33). We constructed a Δ*crtM::kan* mutant because the vectors needed for building transposon mutant libraries do not utilize kanamycin resistance determinants nor is kanamycin resistance routinely used to manipulate the *S. aureus* genome.

SrrAB is a two-component regulatory system utilizing the SrrA DNA-binding response regulator and the membrane-associated histidine kinase SrrB (34, 35). The *srrA* promoter responds to the phosphorylation status of SrrA. We created the pJB38_*srrAp*_*sa*r RBS_*lacZ* plasmid and used it to create a *srrA::lacZ* reporter strain. The expression of beta-galactose from the *saeP1::lacZ* and *srrA::lacZ* reporters and isogenic Δ*crtM::kan* strains were not significantly different (**Figure 4A**). When we serially diluted and spot plated these strains alongside isogenic strains containing *saeR::Tn* or *srrA::Tn*, we noted an increase in visual differentiation in indole formation in the strains lacking staphyloxanthin production (**Figure 4B**). Interestingly, we also observed that strains lacking SrrA displayed sensitivity to X-gal.

**Figure 4.**
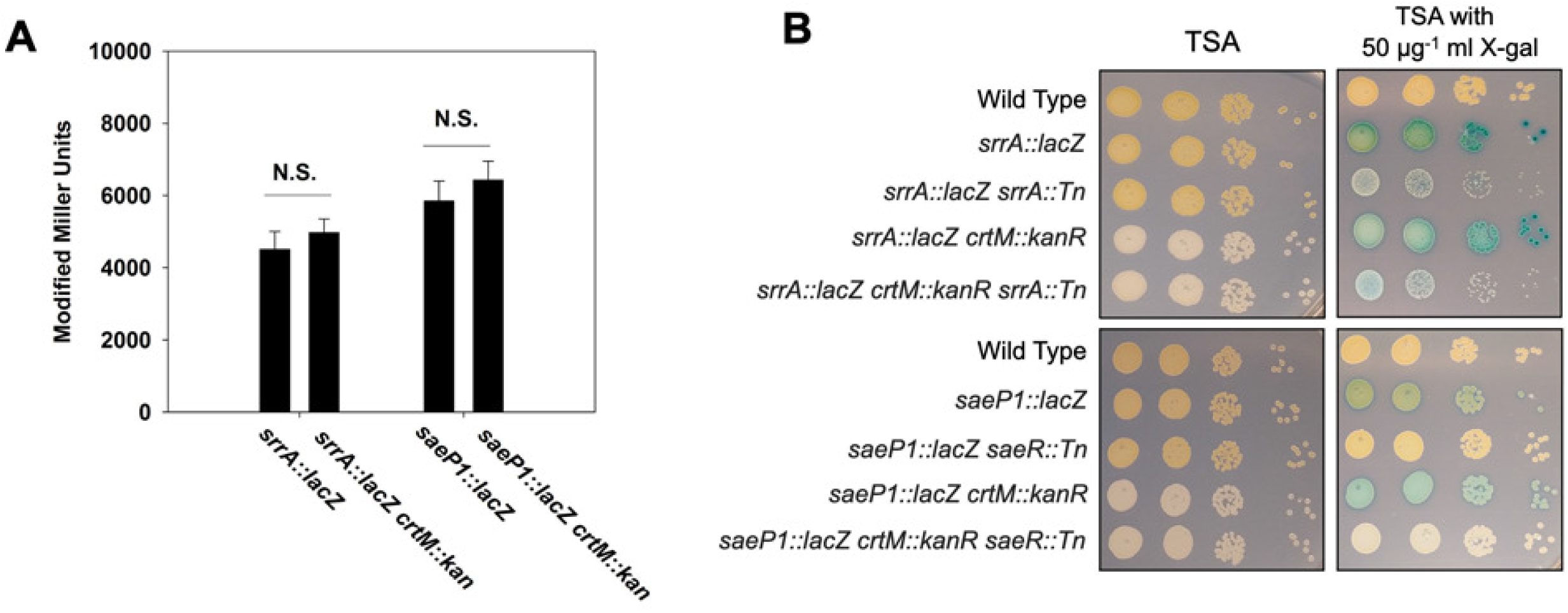
A *crtM::kan* allele prevents staphyloxanthin expression and aids in bluewhite screening. **(A)** β-galactosidase activity in the *saeP1::lacZ* (JMB 9709), *saeP1::lacZ crtM::kanR* (JMB 10021), *srrA::lacZ* (JMB 9742), *srrAp::lacZ crtM::kanR* (JMB 10025) strains during logarithmic or stationary growth phases. Panel B; the following strains were cultured overnight before serial diluting and spot plating on TSA and TSA with 50 μg ml^-1^ X-gal: wild type (JMB 1100), *srrAp::lacZ* (JMB 9742), *srrAp::lacZ srrA::Tn* (JMB 10064), *srrA::lacZ* Δ*crtM::kan* (JMB 10025), *srrA::lacZ* Δ*crtM::kan srrA::Tn* (JMB 10067), *saeP1::lacZ* (JMB 9709), *saeP1::lacZ saeR::Tn* (JMB 9727), *saeP1::lacZ* Δ*crtM::kan* (JMB 10021), *saeP1::lacZ* Δ*crtM::kan saeR::Tn* (JMB 10236). All strains utilized the *sarA* RBS to drive *lacZ* expression. The data displayed in panel A are the average of biological triplicates with the standard deviations shown. Representative images are displayed in panel B.

### Using *lacZ* transcriptional reporter strains for anaerobic screening

Traditionally, *lacZ*-dependent X-gal-hydrolysis screens have only been conducted aerobically because an oxidant (i.e. dioxygen) is necessary for dimerization of the hydrolysis byproduct 5-bromo-4-chloro-3-hydroxyindole, which is visualized as indigo color. *S. aureus* is a facultative anaerobe (36). Blue-white screens have previously been performed after anaerobic growth using X-gal spray overlays (37). We sought to apply this method to visualize β-galactosidase activity in *S. aureus* grown anaerobically on solid medium. We monitored *lacZ* expression in the *srrAp::lacZ* strain, as well as the isogenic *srrB::Tn* and Δ*srrAB::tet* mutants that had been spot plated and cultured anaerobically. The agar plates were colorless after incubation. One plate was sprayed with a 25 mg ml^-1^ X-gal solution and removed from the anaerobic chamber. Another plate also sprayed with a solution of 25 mg ml^-1^ X-gal and 3.3 mg ml^-1^ tetracycline to prevent new β-galactosidase synthesis. Upon exposure to oxygen and indole color development, we observed that the *srrB::Tn* mutant exhibited decreased *srrAp::lacZ* reporter activity compared to the parent strain (**Figure 5A**). The addition of tetracycline had no impact on chromatic development suggesting that the presence of oxygen was not significantly altering *lacZ* expression under the time frame utilized.

**Figure 5.**
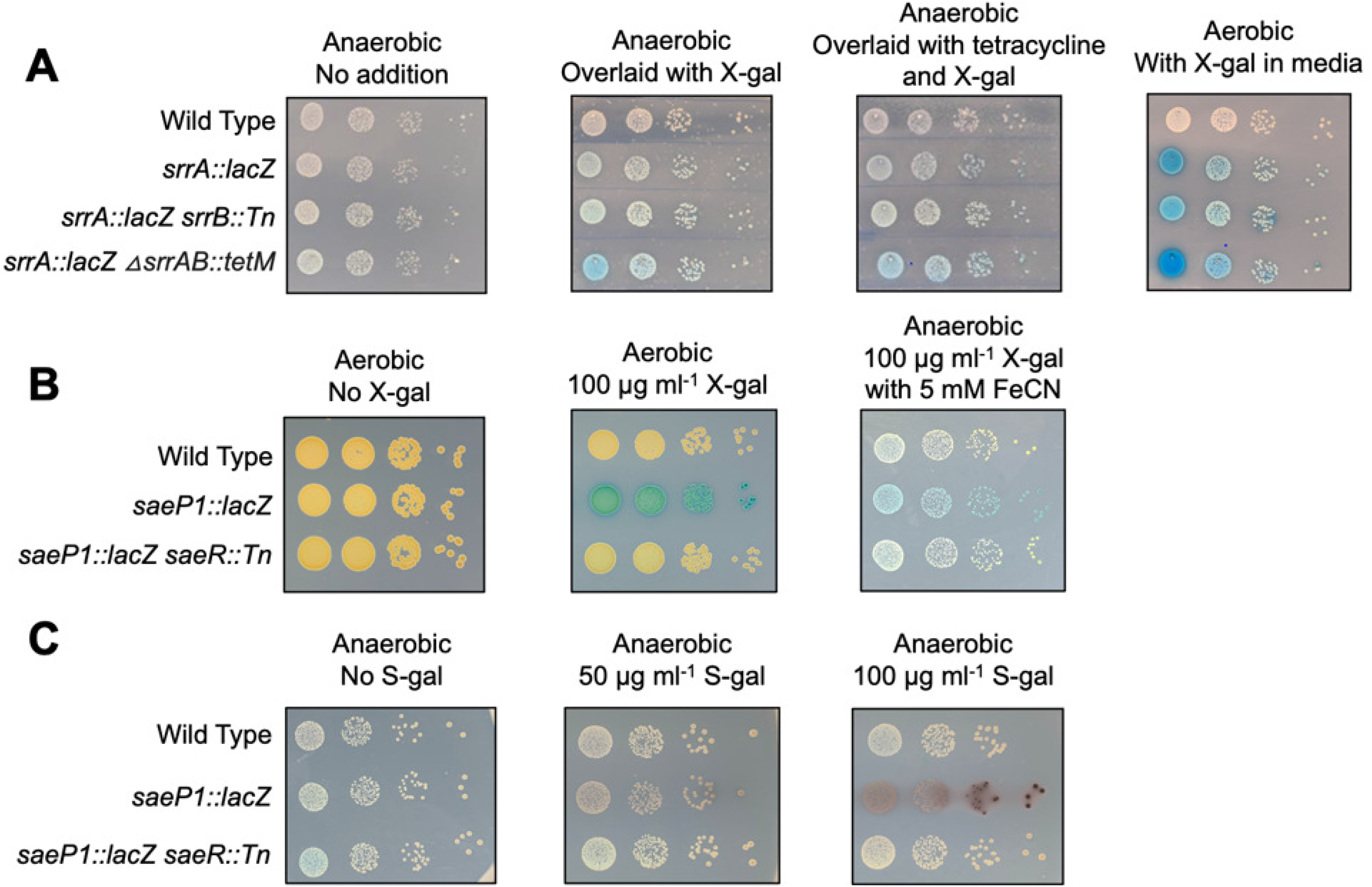
Monitoring *lacZ* expression during in anaerobic growth. **(A)** Overnight cultures of the wild type (JMB 1100), *srrAp::lacZ* (JMB 9742), *srrAp::lacZ srrB::Tn* (JMB 10066), and *srrAp::lacZ* Δ*srrAB::tetM* (JMB 10065) were serial diluted, spotted on TSA, and cultured anaerobically. One plate was sprayed with a 25 mg ml^-1^ solution of X-gal and another was sprayed with a solution of 25 mg ml^-1^ X-gal and 3.3 mg ml^-1^ tetracycline before all three plates were removed from the anaerobic chamber and allowed to develop. **(B)** Overnight cultures of wild type (JMB 1100), *saeP1::lacZ* (JMB 9709), and *saeP1::lacZ saeR::Tn* (JMB 9727) were serial diluted and spotted on TSA with and without 100 mg ml^-1^ X-gal and with and without 5 mM Fe(CN)_6_^3-^ and Fe(CN)_6_^4-^ before incubating in aerobic or anaerobic conditions. **(C)** Overnight cultures of the same strains used in panel B were serial diluted and spotted on plates with or without S-gal. The plates were incubated anaerobically. Representative images are shown.

We next developed a method for monitoring X-gal hydrolysis on solid medium in the absence of dioxygen. The addition of a ferrocyanide-ferricyanide redox system (Fe(CN)_6_^3–/4–^) has been shown to act as an electron acceptor to increase the rate of 5,5’-dibromo-4,4’-dichloro-indigo development when monitoring *lacZ* expression for histochemical analyses (38, 39). We examined whether the addition of these chemicals to solid media would allow us to visualize anaerobic X-gal hydrolysis anaerobically. The inclusion of Fe(CN)_6_^3–/4–^ resulted in the formation of an indigo precipitate anaerobically (**Figure 5B**). Addition of 5 mM Fe(CN)_6_^3–^ and Fe(CN)_6_^4–^ resulted in optimal visualization and little to no growth inhibition. The wild type and the *saeP1::lacZ saeR::Tn* strains did not display noticeable chromophore development, consistent with the blue color arising from *lacZ* expression and X-gal cleavage.

S-gal is a chromogenic substrate for β-galactosidase that can be utilized to examine *lacZ* expression in the absence of oxygen. When hydrolyzed, the byproduct can chelate iron resulting in a black precipitate (40). Importantly, S-gal staining does not require oxygen, and therefore can be utilized to visualize β-galactosidase activity under anaerobic conditions. To visualize anaerobic promoter activity, we spotted *saeP1::lacZ* reporter strains on plates containing S-gal and Fe^3+^ (**Figure 5C**). As expected, the *saeP1::lacZ* strain displayed a black precipitant after anaerobic incubation. The wild type and *saeP1::lacZ saeR::Tn* strains displayed little to no black precipitant suggesting that *lacZ* expression was leading to this phenotype.

### Using a flavin mononucleotide-based reporter system to assay anaerobic transcriptional activity

Flavin mononucleotide-based fluorescent proteins (FbFPs) have been used to develop fluorescent reporter systems that are functional under both aerobic and anaerobic conditions in *E. coli* (12). We generated a multicopy plasmid FbFP-based transcriptional reporter for expression in *S. aureus*. The *E. coli-*derived (*EcFbFP*) genetic sequence was codon-optimized for *S. aureus* expression and placed under the transcriptional control of the *saeP1* promoter. *S. aureus* cells harboring the *saeP1::EcFbFP* reporter displayed significantly greater fluorescence compared to wild-type cells after aerobic culture (**Figure 6A**). Fluorescence was greatly diminished in a Δ*saePQRS::spec* mutant containing *saeP1::EcFbFP* reporter suggesting that the fluorescence signal was specific to *saeP1* promoter activity. Importantly, the *saeP1::EcFbFP* reporter system was functional in the absence of oxygen (**Figure 6B**).

**Figure 6.**
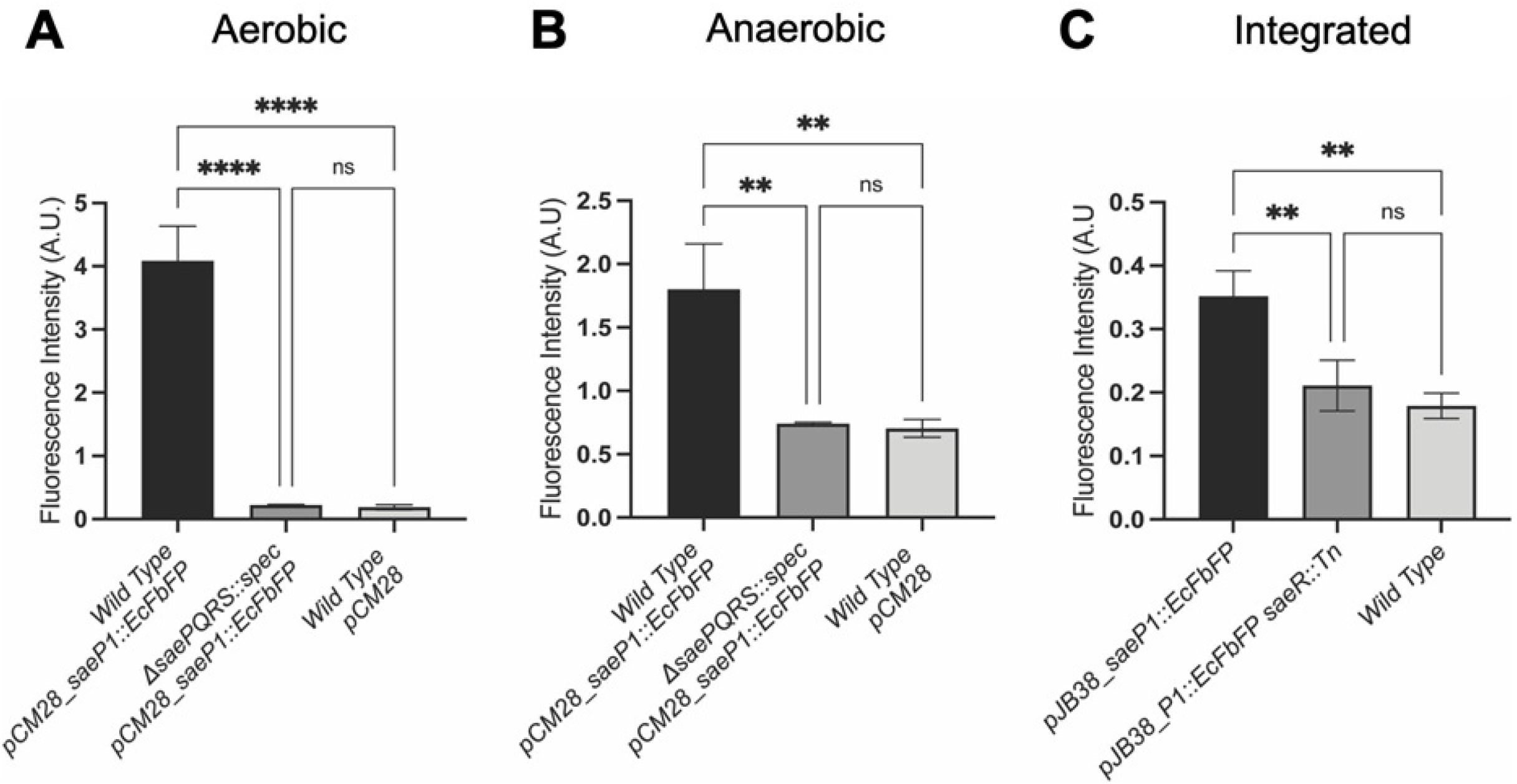
Using *EcFbFP* expression to monitor transcriptional activity. Fluorescence was monitored in the wild type (JMB 1100) and Δ*saePQRS::spec* (JMB 1335) strains containing either pCM28 or pCM28_*saeP1::EcPbFP* after aerobic **(A)** or anaerobic **(B)** culture. **(C)** Fluorescence was monitored in the wild type (JMB 1100), *saeP1::EcFbFP* (JMB 10207), and *saeP1::EcFbFP saeR::Tn* (JMB 10217) strains after aerobic culture. The data represent the average of biological triplicates and standard deviations are shown.

In addition to the plasmid-based *EcFbFP* reporter system, we also generated a chromosomally-integrated *saeP1::EcFbFP* reporter strain, which allows for markerless insertion of the reporter sequence into *geh* of the *S. aureus* genome. The integrated *saeP1::EcFbFP* strain displayed weaker fluorescence signal compared to a strain carrying the plasmid-based reporter (**Figure 6C**). This decrease in signal output likely reflects transcriptional reporter copy number.

### Creation of vectors to utilize hemolysin or lipase expression to monitor transcriptional activity

We generated two vectors that can be used to screen aerobic or anaerobic cultures for altered promoter transcriptional activity using the endogenous *S. aureus* enzymes alpha-hemolysin (*hla*) and lipase (*geh*). Both reporters utilize the *TIR* ribosomal binding site. The vectors provide markerless integration into the *S. aureus* genome allowing for further genetic manipulation. We used the vectors to replace the *hla* or *geh* promoters with the *saeP1* promoter. Notably, the gene products of both *hla* and *geh* are secreted proteins, which allows for screening on solid media.

When examined on rabbit blood agar, *S. aureus* cells containing the *saeP1::hla* reporter produced a zone of hemolysis (**Figure 7A**). Introduction of an *saeP::Tn* or *saeR::Tn* mutation increased and decreased the zone of hemolysis, respectively, suggesting that the changes in *hla* expression were specific to the *saeRS* promoter (**Figure 7B**). Importantly, alpha-hemolysis activity was observed under both aerobic and anaerobic conditions.

**Figure 7.**
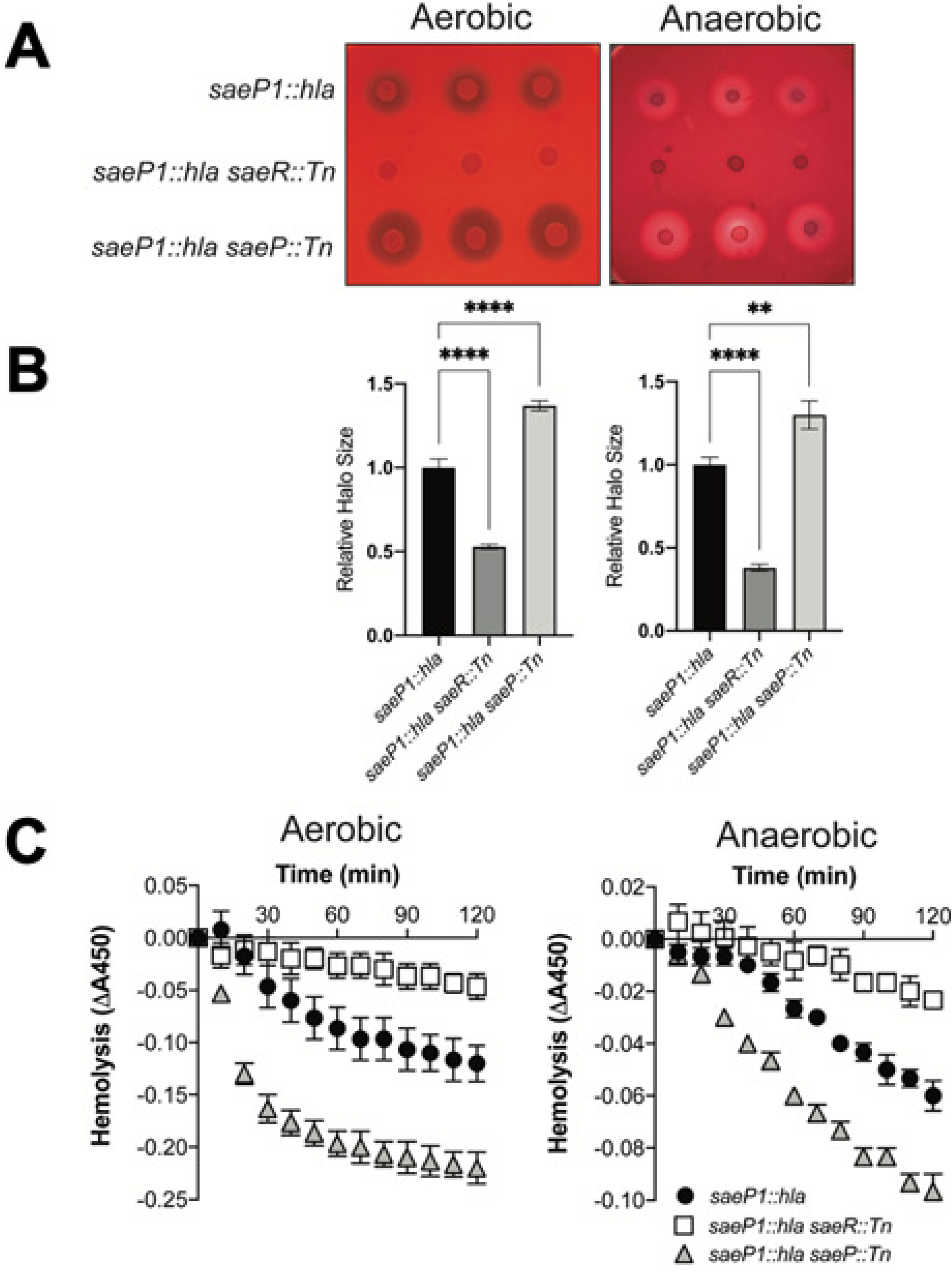
Using *hla* expression to monitor transcriptional activity. **(A)** Overnight cultures of the *saeP1::hla* (JMB 10341), *saeP1::hla saeR::Tn* (JMB 10350), and *saeP1::hla saeP::Tn* (JMB 10351) strains were spotted in triplicate on TSA plates containing 5% defibrinated rabbit blood. **(B)** Quantification of *hla* expression by measuring the clearance zones from the plate image is shown in panel A. Relative halo size was normalized to the average clearance zone of *saeP1::hla* (JMB 10341). **(C)** Quantification of *hla* expression by monitoring hemolysis. Heme content released from rabbit red blood cells was assessed by monitoring absorbance at 450 nm and incubating with spent medium from overnight cultures. The strains utilized are the same as the strains utilized in panel A. The data represent the average of biological triplicates and the error bars represent standard deviations.

To complement the plate-based screening, we quantified alpha-hemolysin activity (**Figure 7C**). We assessed heme content released from rabbit red blood cells incubated with cell-free spent media from overnight cultures. We observed significantly more hemolysis in the spent media from the *saeP::Tn* mutant compared to the parent strain. This result suggests that SaeP plays a role in regulating SaeRS activity in both the presence and absence of oxygen. As expected, we noted significantly decreased hemolysis in the *saeR::Tn* mutant.

We next examined *geh* expression by supplementing the solid medium with Tween-80 substrate and calcium salt, which forms an insoluble precipitate when bound to free fatty acids generated by lipase. *S. aureus* cells harboring the *saeP1::geh* reporter produced a zone of fatty acid precipitate (**Figure 8**). The precipitate zone was significantly smaller and larger upon the introduction of *saeR::Tn* or *saeP::Tn* mutations, respectively. These data suggest that the witnessed *geh* expression was specifically controlled by the transcriptional activity of the *saeP1* promoter. As observed with the alpha-hemolysin reporter system, the *P1::geh* reporter was active under both aerobic and anaerobic conditions.

**Figure 8.**
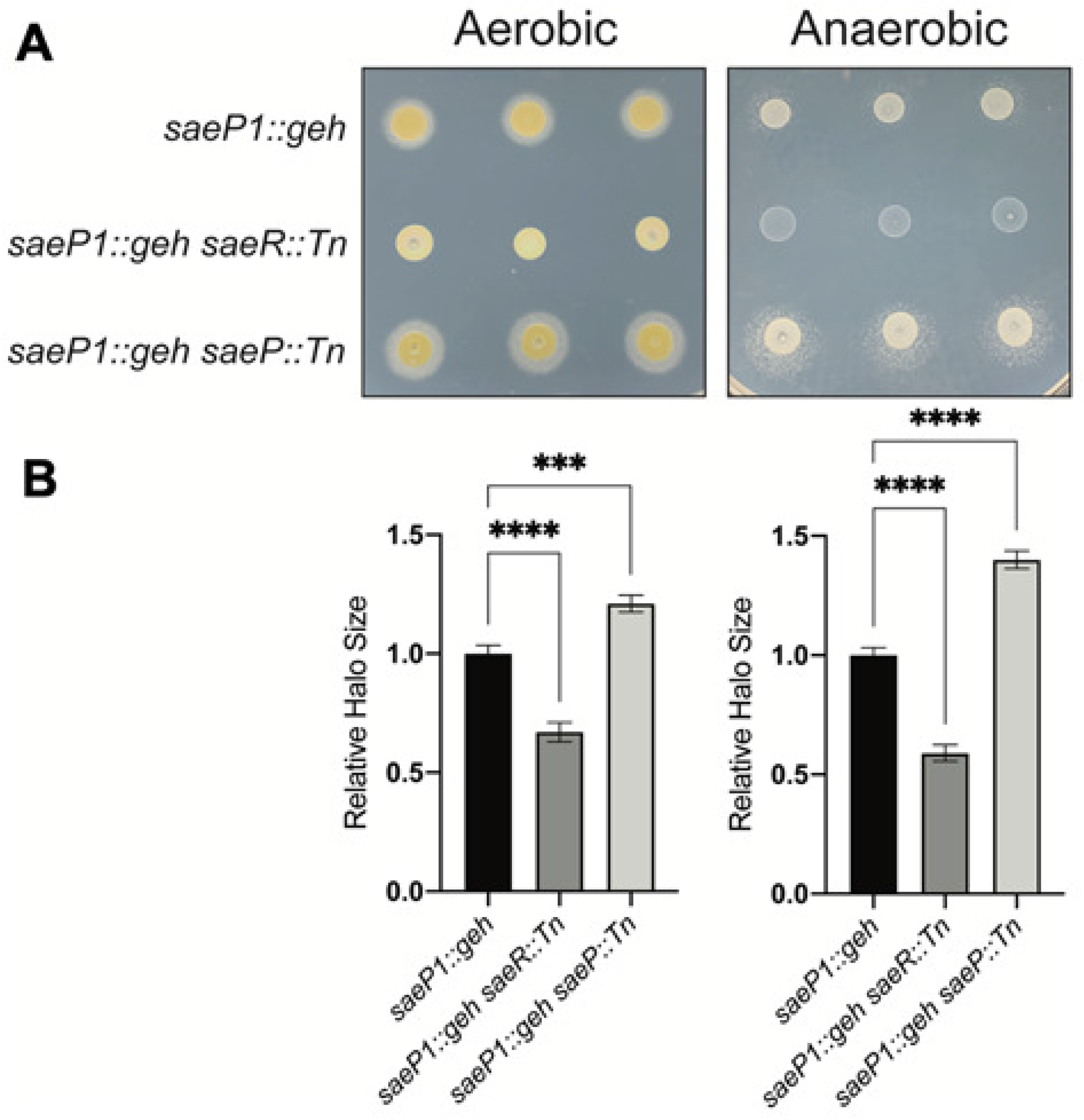
Using *geh* expression to monitor transcriptional activity. **(A)** Overnight cultures of the *saeP1::geh* (JMB 10337), *saeP1::geh saeR::Tn* (JMB 10346), and *saeP1::geh saeP::Tn* (JMB 10347) strains were spotted in triplicate on lipase activity plates containing 1% Tween-80 and calcium salt. **(B)** Quantification of *geh* expression by measuring the precipitate zones from the plate image is shown in panel A. Relative halo size was normalized to the average precipitate zone of *saeP1::geh* (JMB 10337).

## Discussion

We have designed a suite of vectors and methods to monitor transcriptional activity in *S. aureus* under both aerobic and anaerobic growth conditions. These vectors will enable researchers to identify physiological conditions and genetic loci that alter promoter activity. Importantly, these vectors have several key advantages over previously described *S. aureus* vectors. The presence of a yeast cloning cassette and restriction sites that flank the promoter region allow for simple replacement of the promoter of interest using restriction enzyme-based cloning or recombinational cloning. The shuttle vectors can be moved easily between *S. cerevisiae, E. coli*, and *S. aureus*. The vectors have one of four RBS, which allows researchers to tailor reporter gene transcription. There are restriction sites upstream and downstream of the reporter gene allowing for replacement with alternate reporter genes (*gfp, yfp, mCherry*, etc.) Many of the vectors integrate into the non-essential *geh* locus, and to our knowledge, do not hamper the fitness of *S. aureus* under standard laboratory culture conditions. Lastly, the vectors enable researchers to construct markerless reporter strains, which allows for the further genetic manipulation such as the building of transposon mutant libraries using plasmids requiring extensive selection for antibiotic resistance.

*S. aureus* does not show an inherent β-galactosidase activity like other coagulase-positive staphylococci which allows for the monitoring of X-gal hydrolysis [25]. A wide array of β-galactosidase substrates is available for detection of β-galactosidase activity; however, nearly all the substrates described require dioxygen for development. Herein, we demonstrate that anaerobic *S. aureus* gene expression studies can be successfully conducted on solid media with *lacZ-*based reporters using S-gal, an X-gal spray, or the inclusion of an Fe(CN)_6_^3-/4-^ redox cycling system. To our knowledge, this is the first study to report successful usage of an oxidant for monitoring *lacZ* expression during anaerobic growth.

After building transposon mutant libraries in our reporter strains, we found that nearly all the mutants isolated contained mutations that altered staphyloxanthin production, which obfuscated the results from blue-white screening. To circumvent this problem, we created a Δ*crtM::kanR* mutation which prevented staphyloxanthin production and afforded better clarity in plate screening. Kanamycin is rarely used in *S. aureus* molecular biology, and therefore, the Δ*crtM::kanR* mutation allows for further genetic manipulation and for locus movement between strains via transduction.

Three non-*lacZ* based reporter systems allowed us to monitor gene transcriptional activity in the absence of oxygen. Monitoring *saeP1* transcriptional activity by quantifying *EcFbFP* expression worked well when the reporter system was provided as a multi-copy plasmid. The sensitivity of this reporter was diminished when provided in single copy via chromosomal integration even when using the *TIR* RBS to drive expression. The *hla* and *geh* reporters provided robust expression patterns on solid media. Importantly, the activities of both α-hemolysin and lipase do not require oxygen, making these reporter vectors ideal for monitoring anaerobic gene transcription.

As a facultative anaerobe, *S. aureus* can use respiration or fermentation to generate energy and maintain redox homeostasis. Most infection sites are low oxygen or anaerobic environments. Not surprisingly, *S. aureus* alters the transcription of many virulence genes as a variable of oxygen tension. A better understanding of anaerobic gene expression will provide insight into *S. aureus* pathogenesis. The vectors, strains, and methods described herein provide an advanced toolkit to dissect aerobic and anaerobic gene regulation in *S. aureus*.

**Table 1.**
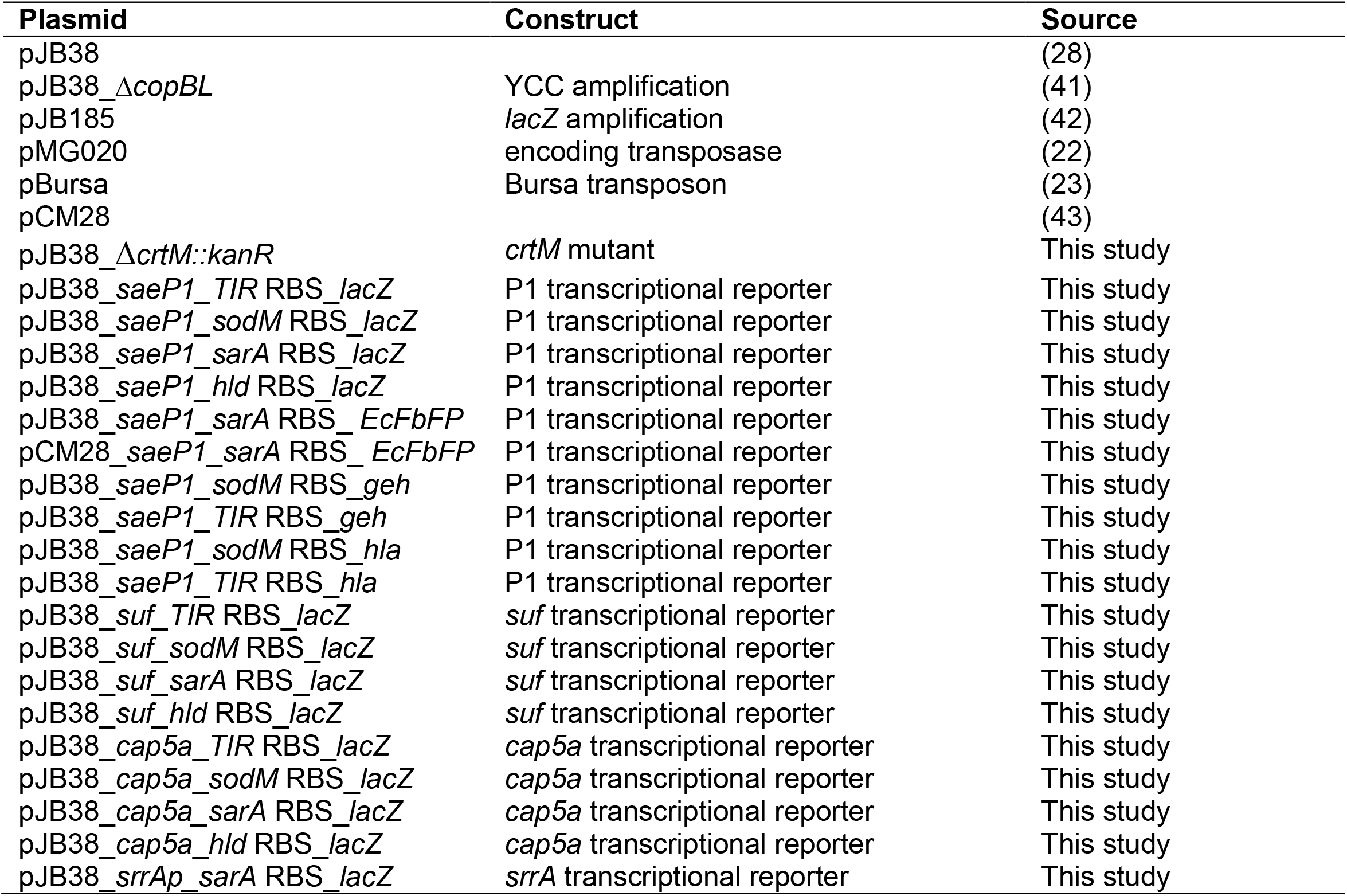
Plasmids utilized in this study.

**Table 2.** *S. aureus* strains utilized in this study.

| **Name** | **Genotype** | **Source** |
| --- | --- | --- |
| RN4220 | Restriction minus | (44) |
| JMB 1100 | Wild type | (43) |
| JMB 2122 | $\Delta bshA::kanR$ | (45) |
| JMB 1335 | $\Delta saePQRS::spec$ | (36) |
| JMB 9741 | *geh::suf_TIR RBS_lacZ* | This study |
| JMB 9739 | *geh::suf_sodM RBS_lacZ* | This study |
| JMB 9740 | *geh::suf_sarA RBS_lacZ* | This study |
| JMB 9738 | *geh::suf_hld RBS_lacZ* | This study |
| JMB 9765 | *geh::cap5a_TIR RBS_lacZ* | This study |
| JMB 9764 | *geh::cap5a_sodM RBS_lacZ* | This study |
| JMB 9754 | *geh::cap5a_sarA RBS_lacZ* | This study |
| JMB 9777 | *geh::cap5a_hld RBS_lacZ* | This study |
| JMB 9709 | *geh::saeP1_sarA RBS_lacZ* | This study |
| JMB 9727 | *geh::saeP1_sarA RBS_lacZ saeR::Tn (ermB)* | This study, (46) |
| JMB 9774 | *geh::saeP1_sarA RBS_lacZ Tn library* | This study |
| JMB 9728 | *geh::saeP1_sarA RBS_lacZ saeP::Tn (ermB)* | This study, (46) |
| JMB 9832 | *geh::saeP1_sarA RBS_lacZ SAUSA300_2501::Tn (crtP) (ermB)* | This study |
| JMB 9964 | *crtM::kanR* | This study |
| JMB 9742 | *geh::srrAp_sarA_RBS_lacZ* | This study |
| JMB 10025 | *geh::srrAp_sarA_RBS_lacZ $\Delta crtM::kanR$* | This study |
| JMB 10021 | *geh::saeP1_sarA RBS_lacZ $\Delta crtM::kanR$* | This study |
| JMB 10067 | *geh::srrAp_sarA_RBS_lacZ $\Delta crtM::kanR$ srrA::Tn (ermB)* | This study, (46) |
| JMB 10236 | *geh::saeP1_sarA RBS_lacZ $\Delta crtM::kanR$ saeR::Tn (ermB)* | This study, (46) |
| JMB 10064 | *geh::srrAp_sarA RBS_lacZ srrA::Tn (ermB)* | This study, (46) |
| JMB 10065 | *geh::srrAp_sarA RBS_lacZ $\Delta srrAB::tetM$* | This study, (34) |
| JMB 10207 | *geh::saeP1_sarA RBS_EcFbFP* | This study |
| JMB 10217 | *geh::saeP1_sarA RBS_EcFbFP saeR::Tn (ermB)* | This study, (46) |
| JMB 10341 | *saeP1_TIR RBS::hla* | This study |
| JMB 10350 | *saeP1_TIR RBS::hla saeR::Tn (ermB)* | This study, (46) |
| JMB 10351 | *saeP1_TIR RBS::hla saeP::Tn (ermB)* | This study, (46) |
| JMB 10337 | *saeP1_TIR RBS::geh* | This study |
| JMB 10346 | *saeP1_TIR RBS::geh saeR::Tn (ermB)* | This study, (46) |
| JMB 10347 | *saeP1_TIR RBS::geh saeP::Tn (ermB)* | This study, (46) |

## References

1. Rohlfing SR, Crawford IP. 1966. Purification and characterization of the beta-galactosidase of Aeromonas formicans. J Bacteriol 91:1085–97.

2. Mandelstam J. 1958. Turnover of protein in growing and non-growing populations of Escherichia coli. Biochem J 69:110–9.

3. Miller JH. 1972. Experiments in molecular genetics, Cold Spring Harbor, NY.

4. Horwitz JP, Chua J, Curby RJ, Tomson AJ, Darooge MA, Fisher BE, Mauricio J, Klundt I. 1964. Substrates for Cytochemical Demonstration of Enzyme Activity. I. Some Substituted 3-Indolyl-Beta-D-Glycopyranosides. J Med Chem 7:574–5.

5. O’Neill AJ, Miller K, Oliva B, Chopra I. 2004. Comparison of assays for detection of agents causing membrane damage in Staphylococcus aureus. J Antimicrob Chemother 54:1127–9.

6. Ranjit DK, Endres JL, Bayles KW. 2011. Staphylococcus aureus CidA and LrgA proteins exhibit holin-like properties. J Bacteriol 193:2468–76.

7. Baum KR, Ahmad Z, Singh VK. 2015. Regulation of Expression of Oxacillin-Inducible Methionine Sulfoxide Reductases in Staphylococcus aureus. Int J Microbiol 2015:617925.

8. Nielsen A, Nielsen KF, Frees D, Larsen TO, Ingmer H. 2010. Method for screening compounds that influence virulence gene expression in Staphylococcus aureus. Antimicrob Agents Chemother 54:509–12.

9. Bojer MS, Baldry M, Ingmer H. 2017. Protocols for Screening Antimicrobial Peptides That Influence Virulence Gene Expression in Staphylococcus aureus. Methods Mol Biol 1548:387–394.

10. Ding Y, Liu X, Chen F, Di H, Xu B, Zhou L, Deng X, Wu M, Yang CG, Lan L. 2014. Metabolic sensor governing bacterial virulence in Staphylococcus aureus. Proc Natl Acad Sci U S A 111:E4981–90.

11. Chapman S, Faulkner C, Kaiserli E, Garcia-Mata C, Savenkov EI, Roberts AG, Oparka KJ, Christie JM. 2008. The photoreversible fluorescent protein iLOV outperforms GFP as a reporter of plant virus infection. Proceedings of the National Academy of Sciences 105:20038–20043.

12. Drepper T, Eggert T, Circolone F, Heck A, Krauß U, Guterl J-K, Wendorff M, Losi A, Gärtner W, Jaeger K-E. 2007. Reporter proteins for in vivo fluorescence without oxygen. Nature Biotechnology 25:443–445.

13. Dinges MM, Orwin PM, Schlievert PM. 2000. Exotoxins of Staphylococcus aureus. Clin Microbiol Rev 13:16-34, table of contents.

14. Rosenstein R, Götz F. 2000. Staphylococcal lipases: Biochemical and molecular characterization. Biochimie 82:1005–1014.

15. Rollof J, Hedstrom SA, Nilsson-Ehle P. 1987. Purification and characterization of a lipase from Staphylococcus aureus. Biochim Biophys Acta 921:364–9.

16. Cadieux B, Vijayakumaran V, Bernards MA, McGavin MJ, Heinrichs DE. 2014. Role of lipase from community-associated methicillin-resistant Staphylococcus aureus strain USA300 in hydrolyzing triglycerides into growth-inhibitory free fatty acids. J Bacteriol 196:4044–56.

17. Novick RP. 1991. Genetic systems in staphylococci. Methods Enzymol 204:587–636.

18. Mashruwala AA, Boyd JM. 2016. De Novo Assembly of Plasmids Using Yeast Recombinational Cloning. Methods Mol Biol 1373:33–41.

19. Joska TM, Mashruwala A, Boyd JM, Belden WJ. 2014. A universal cloning method based on yeast homologous recombination that is simple, efficient, and versatile. J Microbiol Methods 100:46–51.

20. Krute CN, Rice KC, Bose JL. 2017. VfrB Is a Key Activator of the Staphylococcus aureus SaeRS Two-Component System. J Bacteriol 199.

21. Pang YY, Schwartz J, Bloomberg S, Boyd JM, Horswill AR, Nauseef WM. 2013. Methionine Sulfoxide Reductases Protect against Oxidative Stress in Staphylococcus aureus Encountering Exogenous Oxidants and Human Neutrophils. J Innate Immun doi:10.1159/000355915.

22. Grosser MR, Paluscio E, Thurlow LR, Dillon MM, Cooper VS, Kawula TH, Richardson AR. 2018. Genetic requirements for Staphylococcus aureus nitric oxide resistance and virulence. PLoS Pathog 14:e1006907.

23. Bae T, Glass EM, Schneewind O, Missiakas D. 2008. Generating a collection of insertion mutations in the Staphylococcus aureus genome using bursa aurealis. Methods Mol Biol 416:103–16.

24. Bose JL, Daly SM, Hall PR, Bayles KW. 2014. Identification of the Staphylococcus aureus vfrAB operon, a novel virulence factor regulatory locus. Infect Immun 82:1813–22.

25. Luong TT, Lee CY. 2007. Improved single-copy integration vectors for Staphylococcus aureus. J Microbiol Methods 70:186–90.

26. Malone CL, Boles BR, Lauderdale KJ, Thoendel M, Kavanaugh JS, Horswill AR. 2009. Fluorescent reporters for Staphylococcus aureus. J Microbiol Methods 77:251–60.

27. Bose JL. 2014. Genetic manipulation of staphylococci. Methods Mol Biol 1106:101–11.

28. Bose JL, Fey PD, Bayles KW. 2013. Genetic tools to enhance the study of gene function and regulation in Staphylococcus aureus. Appl Environ Microbiol 79:2218–24.

29. Roberts CA, Al-Tameemi HM, Mashruwala AA, Rosario-Cruz Z, Chauhan U, Sause WE, Torres VJ, Belden WJ, Boyd JM. 2017. The Suf Iron-Sulfur Cluster Biosynthetic System Is Essential in Staphylococcus aureus, and Decreased Suf Function Results in Global Metabolic Defects and Reduced Survival in Human Neutrophils. Infect Immun 85.

30. Mashruwala AA, Gries CM, Scherr TD, Kielian T, Boyd JM. 2017. SaeRS Is Responsive to Cellular Respiratory Status and Regulates Fermentative Biofilm Formation in Staphylococcus aureus. Infect Immun 85.

31. Jeong DW, Cho H, Jones MB, Shatzkes K, Sun F, Ji Q, Liu Q, Peterson SN, He C, Bae T. 2012. The auxiliary protein complex SaePQ activates the phosphatase activity of sensor kinase SaeS in the SaeRS two-component system of Staphylococcus aureus. Mol Microbiol 86:331–48.

32. Liu CI, Liu GY, Song Y, Yin F, Hensler ME, Jeng WY, Nizet V, Wang AH, Oldfield E. 2008. A cholesterol biosynthesis inhibitor blocks Staphylococcus aureus virulence. Science 319:1391–4.

33. Pelz A, Wieland KP, Putzbach K, Hentschel P, Albert K, Gotz F. 2005. Structure and biosynthesis of staphyloxanthin from Staphylococcus aureus. J Biol Chem 280:32493–8.

34. Mashruwala AA, Boyd JM. 2017. The Staphylococcus aureus SrrAB Regulatory System Modulates Hydrogen Peroxide Resistance Factors, Which Imparts Protection to Aconitase during Aerobic Growth. PLoS One 12:e0170283.

35. Yarwood JM, McCormick JK, Schlievert PM. 2001. Identification of a novel two-component regulatory system that acts in global regulation of virulence factors of Staphylococcus aureus. J Bacteriol 183:1113–23.

36. Mashruwala AA, Van De Guchte A, Boyd JM. 2017. Impaired respiration elicits SrrAB-dependent programmed cell lysis and biofilm formation in Staphylococcus aureus. eLife 6.

37. Lie TJ, Leigh JA. 2007. Genetic screen for regulatory mutations in Methanococcus maripaludis and its use in identification of induction-deficient mutants of the euryarchaeal repressor NrpR. Appl Environ Microbiol 73:6595–600.

38. Lojda Z. 1970. Indigogenic methods for glycosidases. II. An improved method for beta-D-galactosidase and its application to localization studies of the enzymes in the intestine and in other tissues. Histochemie 23:266–88.

39. Trifonov S, Yamashita Y, Kase M, Maruyama M, Sugimoto T. 2016. Overview and assessment of the histochemical methods and reagents for the detection of beta-galactosidase activity in transgenic animals. Anat Sci Int 91:56–67.

40. Heuermann K, Cosgrove J. 2001. S-Gal(tm): An Autoclavable Dye for Color Selection of Cloned DNA Inserts. BioTechniques 30:1142–1147.

41. Al-Tameemi H, Beavers WN, Norambuena J, Skaar EP, Boyd JM. 2020. Staphylococcus aureus lacking a functional MntABC manganese import system has increased resistance to copper. Mol Microbiol doi:10.1111/mmi.14623.

42. Austin CM, Garabaglu S, Krute CN, Ridder MJ, Seawell NA, Markiewicz MA, Boyd JM, Bose JL. 2019. Contribution of YjbIH to Virulence Factor Expression and Host Colonization in Staphylococcus aureus. Infect Immun 87.

43. Mashruwala AA, Pang YY, Rosario-Cruz Z, Chahal HK, Benson MA, Mike LA, Skaar EP, Torres VJ, Nauseef WM, Boyd JM. 2015. Nfu facilitates the maturation of iron-sulfur proteins and participates in virulence in Staphylococcus aureus. Mol Microbiol 95:383–409.

44. Kreiswirth BN, Lofdahl S, Betley MJ, O’Reilly M, Schlievert PM, Bergdoll MS, Novick RP. 1983. The toxic shock syndrome exotoxin structural gene is not detectably transmitted by a prophage. Nature 305:709–12.

45. Rosario-Cruz Z, Chahal HK, Mike LA, Skaar EP, Boyd JM. 2015. Bacillithiol has a role in Fe-S cluster biogenesis in Staphylococcus aureus. Mol Microbiol 98:218–42.

46. Fey PD, Endres JL, Yajjala VK, Widhelm TJ, Boissy RJ, Bose JL, Bayles KW. 2013. A genetic resource for rapid and comprehensive phenotype screening of nonessential Staphylococcus aureus genes. MBio 4:e00537–12.

